# The Distribution of Nitric Oxide-Synthesizing Neurons and Soluble Guanylate Cyclase in the Pigeon Brain

**DOI:** 10.1101/2025.03.24.644994

**Authors:** Alina Steinemer, Marie Ziegler, Kevin Haselhuhn, Onur Güntürkün, Noemi Rook

## Abstract

Nitric oxide (NO) is a diffusible neuromodulator with roles in synaptic plasticity and memory flexibility, exerting its primary effects via the enzyme soluble guanylate cyclase (sGC). Despite its well-documented functions in mammals and insects, little is known about the neuroanatomical distribution and functional relevance of NO in birds, particularly in relation to dopaminergic systems. This study used histochemical and immunohistochemical techniques to map the distribution of NO-synthesizing neurons—identified by NADPH-diaphorase (NADPH-d) and nNOS activity—and their relation to sGC and tyrosine hydroxylase (TH)-positive dopaminergic pathways in the pigeon brain. We found extensive NADPH-d labeling throughout forebrain, midbrain, and hindbrain regions. Among TH-positive midbrain structures, the locus coeruleus exhibited high colocalization with nNOS, while moderate colocalization was seen in the ventral tegmental area substantia grisea centralis and substantia nigra. Notably, a significant proportion of sGC-positive neurons was targeted by TH and NADPH-d positive fibres in the pigeon NCL. Our findings support the potential for NO-dopamine interactions in avian species, reminiscent of memory-related mechanisms in *Drosophila melanogaster*, and contribute to an understanding of conserved pathways that may underlie flexible learning and memory processing during navigation or related tasks across vertebrates. This work also offers insight into comparative NADPH-d distribution among avian species, with implications for aging, spatial learning, and memory formation.

## Introduction

In ever-changing environments, animals must adapt their memories to ensure survival, employing mechanisms such as extinction learning (Packheiser et al., 2021). Recent discoveries in *Drosophila melanogaster* have revealed an elegant mechanism supporting such memory flexibility: a subset of dopaminergic neurons releases both dopamine and nitric oxide (NO), enabling the fly to both reinforce and gradually “forget” memories associated with rewards or punishments (Aso et al., 2019; Green & Lin, 2020). In this system, dopamine strengthens memory traces tied to meaningful experiences, while NO serves as a complementary, slowly acting signal that weakens these memories over time. This combination of reinforcement and decay ensures that memories remain adaptable—an essential trait for navigating dynamic surroundings. Interestingly, some neurons release dopamine but not NO; creating a heterogeneous ensemble of neurons that keep up the trace for different lengths of time.

The striking role of NO in memory adaptability in flies raises an intriguing question: could vertebrate brains also rely on NO for similar purposes? In mammals, NO is known to influence synaptic plasticity, learning, memory formation, neurogenesis, and neuroprotection (Bon & Garthwaite, 2003; Zoubovsky et al., 2011). NO is produced from L-arginine by neuronal nitric oxide synthase (nNOS) in a process requiring reduced nicotinamide adenine dinucleotide phosphate (NADPH) as a cofactor (Förstermann & Sessa, 2012). As a diffusible gas, NO acts on various molecular targets, with its most significant effects mediated through soluble guanylate cyclase (sGC), which transduces NO signals to downstream pathways involved in neuronal function (Krumenacker et al., 2004).

Most knowledge of NO distribution in the brain comes from studies on human and rodent tissue, leaving NO distribution in birds less well understood. Nonetheless, several avian species have been investigated for NO-synthesizing neurons, including Japanese quail (*Coturnix japonica*; (Panzica et al., 1994)), rock pigeon (*Columba livia*; (Atoji et al., 2001)), budgerigar (*Melopsittacus undulates*; (Cozzi et al., 1997)), and chicken (*Gallus domesticus*; (Brüning, 1993)). Across these studies, NO-synthesizing neurons were found in a variety of telencephalic, thalamic, midbrain, hindbrain, and cerebellar areas. However, species differences were noted, particularly in regions like the hippocampus and olfactory bulb (Atoji et al., 2001). In zebra finches (*Taeniopygia guttata*), investigations focused specifically on NO-synthesizing neurons within the song system (Wallhäusser-Franke et al., 1995), indicating specialized roles in vocal learning.

In contrast to the relatively well-documented NO synthesis, detailed information on the distribution of NO receptor molecules, such as soluble guanylate cyclase (sGC), is only available for rodents (Ariano et al., 1982; Ding et al., 2004; Furuyama et al., 1993) and has not yet been explored in any avian species. This gap in avian research also leaves us without insights into the potential colocalization of NO-synthesizing neurons with sGC, although such colocalization has been observed in rats (Ding et al., 2004; Schmidt et al., 1992).

Furthermore, evidence from rodents indicates significant interaction between NO and dopamine systems. For example, a notable number of neurons in the rat ventral tegmental area co-express both NO and dopamine (Klejbor et al., 2004), a pattern similarly found in the human temporal cortex (Benavides-Piccione & DeFelipe, 2003). These findings align with earlier studies suggesting NO’s role in modulating dopamine release and uptake (Kiss et al., 1999; Pogun et al., 1994). Although comparable studies in birds have not yet been conducted, a similar NO-dopamine relationship may exist within avian species as well. This lack of information leaves an open question: could birds also exhibit a NO-mediated mechanism for memory flexibility akin to that in fruit flies?

To address this, the present study investigates the distribution of NO-synthesizing neurons in the pigeon brain, focusing on their colocalization with tyrosine hydroxylase (TH), an enzyme critical in dopamine synthesis, as well as with sGC. By exploring whether these NO pathways overlap with dopaminergic systems, we aim to determine if a similar neuroanatomical framework for adaptive memory processing is present in avian species.

Due to its gaseous nature, NO cannot be visualized directly; therefore, we employed immunohistochemical staining of nNOS and histochemical staining of NADPH-diaphorase (NADPH-d), widely used markers for NO-synthesizing neurons (Suárez et al., 2006). This combined approach not only clarifies NO’s anatomical distribution in the pigeon brain but also allows us to assess whether NO-dopamine interactions that support memory flexibility in fruit flies might have an evolutionary basis in vertebrates as well. Findings from this study could provide significant insights into conserved mechanisms of memory dynamics across species, offering a broader understanding of how neural systems support flexible memory.

## Methods

### Experimental subjects

This study included a total of 16 homing pigeons *(Columba livia),* from which four were used for a combined staining of TH and NADPH-d, two for a triple staining of NADPH-d, TH and sGC, six for a combined NOS and TH staining, and four for a single NADPH-d staining. In accordance with the 3R principles of ethical animal research—Replacement, Reduction, and Refinement—this study was conducted using brain tissue from pigeons that had been perfused as part of unrelated, ethically approved experiments. No animals were sacrificed specifically for the purposes of this study. All animals were obtained from local breeders and were individually housed in wire-mesh cages. Housing rooms were controlled for temperature, humidity, and day/night cycles (12-h light–dark cycle). All experiments were performed according to the guidelines regarding the care and use of animals adopted by the German Animal Welfare Law for the prevention of cruelty to animals in agreement with the Directive 2010/63/EU of the European Parliament and of the Council of 22 September 2010, and were approved by the animal ethics committee of the Landesamt für Natur, Umwelt und Verbraucherschutz NRW, Germany. All efforts were made to minimize the number of animals used and to minimize their suffering.

### Perfusion and tissue processing

For perfusion, animals received an intramuscular injection of 0.1 ml heparin (B. Braun, Melsungen, Germany) diluted in 0.1 ml of 0.9 % sodium chloride (NaCl) to prevent blood clots during perfusion. After a dwell time of 10 min, equithesin (0.45 ml/100g body weight) was injected either intramuscularly into the breast muscle or intravenously into the brachial vein. As soon as both eyelid closure and toe pinching reflexes were negative and the heart stopped beating, animals were transcardially perfused with 500 ml of 0.9 % NaCl (40°C) followed by ice-cold (4 °C) 4% paraformaldehyde (PFA) in 0.12 M phosphate-buffered saline (PBS; pH 7.4). After that, the fixated brains were dissected and incubated in a postfix solution (30 % sucrose in 4% PFA) for 2 h at 4°C. For cryoprotection, brains were transferred into sucrose solution (30% sucrose in PBS) for 24 h. Subsequently, brains were embedded in 15% gelatin dissolved in 30% sucrose solution for optimal stereotactic alignment, and again fixated in 4% PFA solution for 24 h. Embedded brains were then transferred into 30% sucrose solution for at least 24 h. Finally, brains were cut into 40 µm thick slices in frontal plane using a rotating microtome (Leica, Wetzlar, Germany). Slices were collected in ten series per bird and stored in storage buffer (0.1% sodium azide in PBS) at 4°C until further processing.

### Histochemistry and immunohistochemistry

For all stainings, every tenth slice of a brain was used and all procedures were conducted with free floating slices. First, we aimed to visualize TH, a precursor of dopamine, in combination with NADPH-d activity, which is colocalized with nNOS in brain tissue (Dawson et al., 1991; Hope et al., 1991; Saffrey et al., 1992; Tracey et al., 1993; Young et al., 1997). We conducted the DAB reaction first to visualize TH, followed by an incubation in nitroblue tetrazolium and NADPH to visualize NADPH-d activity (Wallhäusser-Franke et al., 1995). Slices were first rinsed in PBS (3 x 10 min), and then incubated in 0.3% hydrogen peroxide (H_2_O_2_) in distilled water for 30 min to block endogenous peroxidases. After that, slices were rinsed again (3 x 10 min in PBS) and incubated in 10% normal horse serum (Vectastain Elite ABC kit, Vector Laboratories, Newark, USA) in PBS containing 0.3% Triton-X-100 (PBST) for 60 min to block unspecific binding sites. Slices were transferred to the primary antibody solution containing a mouse anti-TH antibody (1:1000 in PBST; Calbiochem, San Diego, USA) overnight at 4°C. On the next day, slices were again rinsed (3 x 10 min in PBS) and then incubated in the secondary antibody solution containing a biotinylated goat anti-mouse antibody (1:500 in PBST; Vectastain Elite ABC kit, Vector Laboratories, Newark, USA) for 60 min at room temperature. Subsequently, slices were rinsed (3 x 10 min in PBS) and transferred to an avidin-biotin-peroxidase solution (1:100 in PBST each; Vectastain Elite ABC kit, Vector Laboratories, Newark, USA) for 60 min. After further rinsing (3 x 10 min in PBS followed by 1 x 5 min in 0.1 M sodium acetate buffer, pH 6.0), slices were incubated in DAB solution which was composed of DAB (0.2 mg/ml), cobalt chloride (0.4 mg/ml), ammonium chloride (0.4 mg/ml), and β-D-glucose (4 mg/ml) in 0.1 M sodium acetate buffer. We did not add ammonium nickel sulfate here in order to obtain a brown instead of a black reaction product that could be differentiated more easily from the following nitroblue NADPH-d staining. The DAB reaction was initiated by adding glucose oxidase (80−100 µl/50 ml DAB solution) which led to the oxidation of the β-D-glucose. During the reaction, staining intensity was visually controlled continuously, and the solution was changed every 10 min. The staining reaction was stopped after 30 min by rinsing the slices in 0.1 M sodium acetate buffer (3 x 5 min) and PBS (3 x 5 min). For the following NADPH-d staining, slices were incubated in a solution containing 1 mM NADPH (Sigma-Aldrich, St. Louis, USA) and 0.5 mM nitroblue tetrazolium (Sigma-Aldrich, St. Louis, USA) diluted in PBS containing 2% Triton-X-100 for 60 – 90 min. The incubation was conducted in the dark at 37°C. With the same NADPH staining protocol, another set of brain sections was exclusively stained for NADPH. Finally, slices were rinsed (5 x 10 min in PBS) and mounted onto gelatin-coated slides. They were dehydrated in ethanol and xylene, and cover slipped with DPX (Fluka, Munich, Germany).

Next, we conducted fluorescent stainings to search for possible colocalization of TH and nNOS. For that, slices were first rinsed in PBS (3 x 10 min) and then incubated in 10% normal horse serum (Vectastain Elite ABC kit, Vector Laboratories, Newark, USA) in PBST for 60 min to block unspecific binding sites. Without rinsing, slices were transferred to the primary antibody solution. The solution contained a polyclonal rabbit anti-TH antibody (Sigma-Aldrich, St. Louis, USA, 1:2000) and a polyclonal goat anti-nNOS antibody (Invitrogen, Darmstadt, Germany, 1:1000) diluted in PBST with 5% bovine serum albumin. After 72 hours of incubation at 4°C, slices were rinsed in PBS (3 x 10 min) and transferred to the secondary antibody solution containing donkey anti-rabbit AlexaFluor405 (Invitrogen, Darmstadt, Germany, 1:500) and donkey anti-goat AlexaFluor594 (Invitrogen, Darmstadt, Germany, 1:500) diluted in PBST for 60 min at room temperature. After final rinsing in PBS (3 x 10 min), slices were mounted onto glass slides (Superfrost Plus, Thermo Scientific) and embedded with Fluoromount-G (SouthernBiotech, Birmingham, USA). We ensured that exposure to light was reduced to a minimum in order to preserve as much fluorescence as possible.

Moreover, we conducted a triple staining against sGC, NADPH and TH. Therefore, we started with a DAB staining against sGC, followed by a NADPH staining as described above and finished with a fluorescence staining against TH. More specifically, for the GC staining free floating slices were stained permanently by conducting a DAB (3,3-diaminobenzidinetetrahydrochloride) reaction with a commercially available DAB-Kit (DAB Substrate Kit SK-4100, Vector Laboratories, Newark, USA). Slices were first rinsed in PBS (pH 7.4; 3 x 10 min). Then they were incubated in 0.3% hydrogen peroxide (H2O2) in distilled water for 30 min to block endogenous peroxidases. After that, slices were again rinsed in PBS (3 x 10 min) and incubated in 10% normal horse serum (Vectastain Elite ABC kit, Vector Laboratories, Newark, USA) in PBS containing 0.3% Triton-X-100 (PBST) for 60 min to block unspecific binding sites. Subsequently, slices were incubated in a polyclonal rabbit anti-GC (FabGennix, Frisco, USA, 1:1000 in PBST) for 72 hours at 4°C. After that, slices were rinsed in PBS (3 x 10 min) and incubated in a secondary biotinylated donkey anti-rabbit antibody (Jackson ImmunoResearch, Ely, UK, 1:500) diluted in PBST for 60 min at room temperature. After further rinsing in PBS (3 x 10 min), they were incubated in an avidin-biotin complex solution (Vectastain Elite ABC kit, Vector Laboratories, Newark, USA) diluted in PBST (1:100 each) for 60 min. After that, slices were rinsed in PBS (3 x 10 min) and transferred to the DAB working solution containing 2 drops (50 μl) of DAB stock solution, 1 drop (42 μl) of buffer stock solution, and 1 drop (40 μl) of nickel solution per 5 ml distilled water. The staining reaction was carried out in cell wells and each well contained 2 ml of DAB working solution. To initiate the reaction, 3 µl H2O2 were added to each well and after 90 s incubation time, slices were transferred to PBS to stop the reaction. They were then rinsed in PBS (3 x 10 min) for the last time. For the TH fluorescence staining, slices were first rinsed in PBS (3 x 10 min) and then incubated in 10% normal horse serum (Vectastain Elite ABC kit, Vector Laboratories, Newark, USA) in PBST for 60 min to block unspecific binding sites. Without rinsing, slices were transferred to the primary antibody solution containing a polyclonal rabbit anti-TH antibody (Sigma-Aldrich, St. Louis, USA, 1:500) diluted in PBST with 5% bovine serum albumin. After 72 hours of incubation at 4°C, slices were rinsed in PBS (3 x 10 min) and transferred to the secondary antibody solution containing donkey anti-rabbit AlexaFluor594 (Invitrogen, Darmstadt, Germany, 1:200) diluted in PBST for 60 min at room temperature. After final rinsing in PBS (3 x 10 min), slices were mounted onto glass slides (Superfrost Plus, Thermo Scientific) and embedded with Fluoromount-G (SouthernBiotech, Birmingham, USA). We ensured that exposure to light was reduced to a minimum in order to preserve as much fluorescence as possible.

### Microscopic analysis

All slices were imaged bilaterally at 100x magnification with a ZEISS Axio Scan.Z1 (9.94x/0.45 Hamamatsu Orca Flash). Images were then processed using ZEN blue 3.7 (ZEISS, Jena, Germany).

## Results

### Distribution of nitric oxide-synthesizing neurons and tyrosine hydroxylase

In order to investigate the potential role of nitric oxide (NO) in memory flexibility within the pigeon brain, we assessed the distribution of NO-synthesizing neurons alongside dopaminergic projections, visualized by nicotinamide adenine dinucleotide phosphate (NADPH) diaphorase (NADPH-d) and tyrosine hydroxylase (TH) staining (Figure 1). The distinct labeling of NADPH-d for NO-synthesizing neurons and TH for dopamine pathways allows us to observe their specific and overlapping patterns across brain regions. We utilized brain tissue from four pigeons, applying NADPH-d and TH staining to the same slices. This dual-staining approach revealed four prominent types of NADPH-d labeling (Figure 1). We saw subtle, widespread capillary staining; (Figure 1 A, B) dense neuropil staining, often regional; (Figure 1 C) unspecific background staining, which was useful for delineating borders between nuclei (Figure 1 D); and clear visualization of individual neurons, with distinct dendritic and axonal projections (Figure 1 E, F). Most NADPH-d-positive neurons were easily distinguishable from TH-positive structures, facilitating a thorough mapping of each staining type across various brain regions. All coordinates were obtained from the pigeon brain atlas by Karten and Hodos (1967).

**Figure 1:**
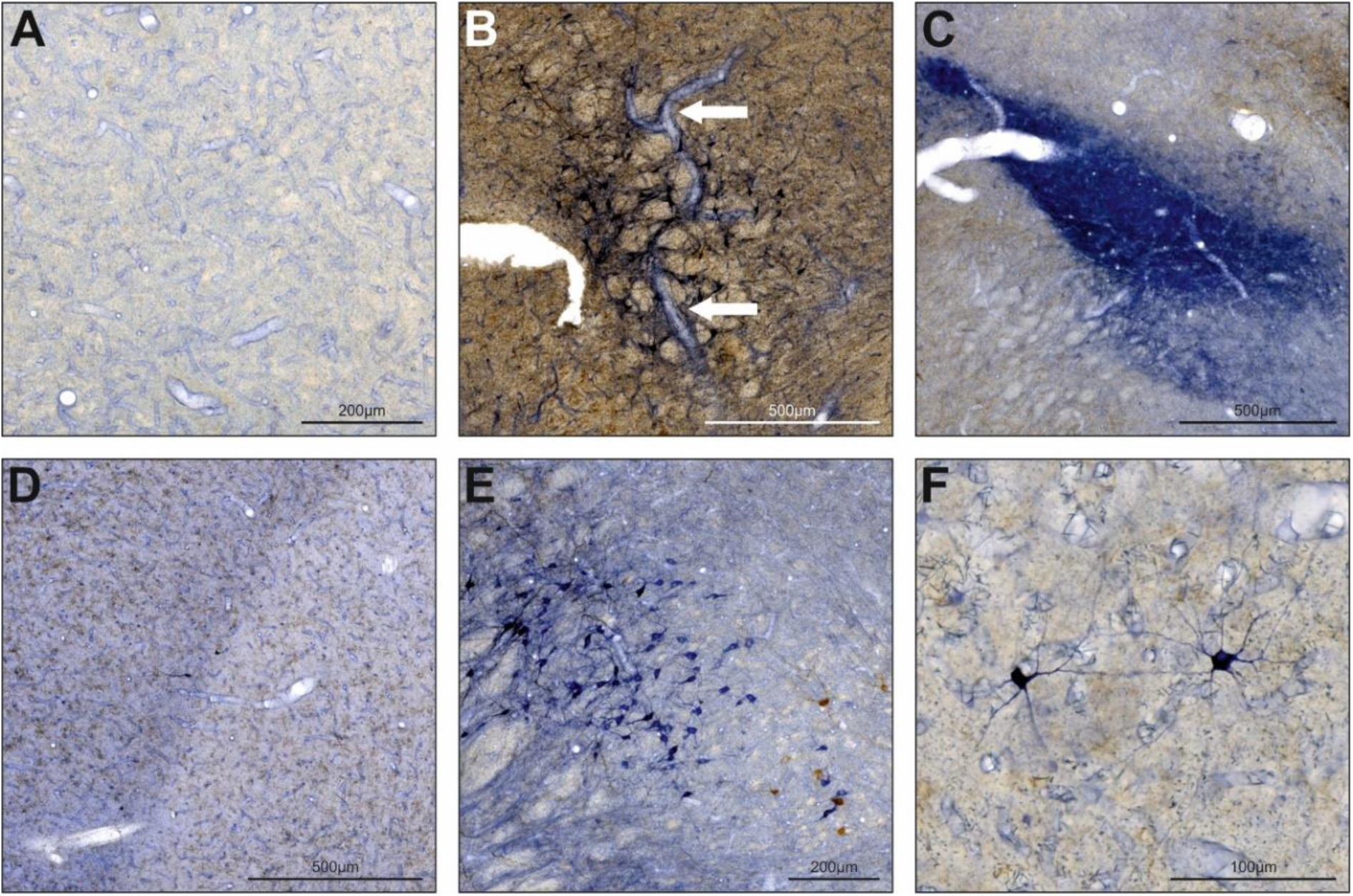
Reaction products following NADPH-d staining. **(A)** Comprehensive staining of capillaries. The exemplary picture displays Rt. **(B)** Rare staining of prominent blood vessels. The white arrows indicate a vessel in the midbrain. The brown background reflects a dense dopaminergic innervation of this area. **(C)** Intensive neuropil staining. The exemplary picture displays SpM. **(D)** Unspecific background staining. Together with staining of capillaries, this was observed throughout the whole brain. Here, the border between HA and HI is clearly visible. **(E)** and **(F)** Individually stained neurons. Somata, dendrites, and axons were usually easy to distinguish from background and TH-positive structures which were stained in brown. For abbreviations, see list.

We found NADPH-d-positive neurons, fibers, or neuropil exist in almost every region of the pigeon brain, indicating a very versatile and/or generalized role of NO. The most prominently labeled regions included OB, CPP, TPO, TuO, LFB, POA, SM, HM, RS, LA, GLv, GLdp, SpM, LHy, IS, ICo, MLd, nIV, MPv, IP, PM, PL, LLd, CS, nBOR, and certain layers of TeO (Figure 2). In general, NADPH-d activity was least intense in endbrain regions and increased towards thalamic and midbrain regions (Figure 2). However, compared to previous NADPH-d studies labeling in the entopallium, arcopallium, TnA, Rt, VTA, Ru, Imc, Ipc, and VLV was weaker than expected (Figure 2). Within TeO, we detected three bands of positive neurons. Although we investigated, TH and NADPH distributions within the whole brain (Figure 2), our result section will especially focus on the telencephalon, which is the major target of TH positive fibres and relevant TH positive midbrain structures to assess the neuroanatomical framework of the memory flexibility theory that has been established in fruit flies.

**Figure 2:**
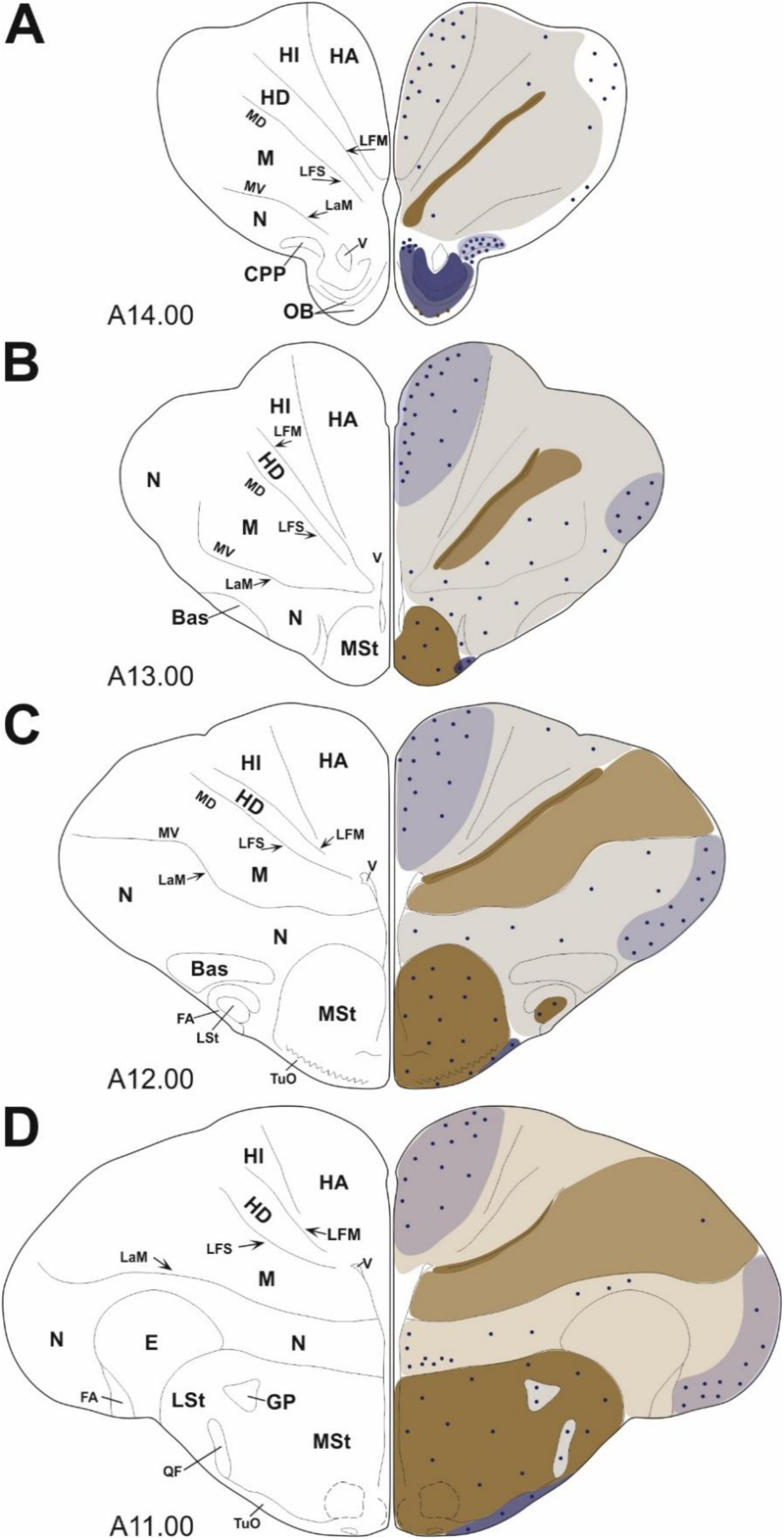
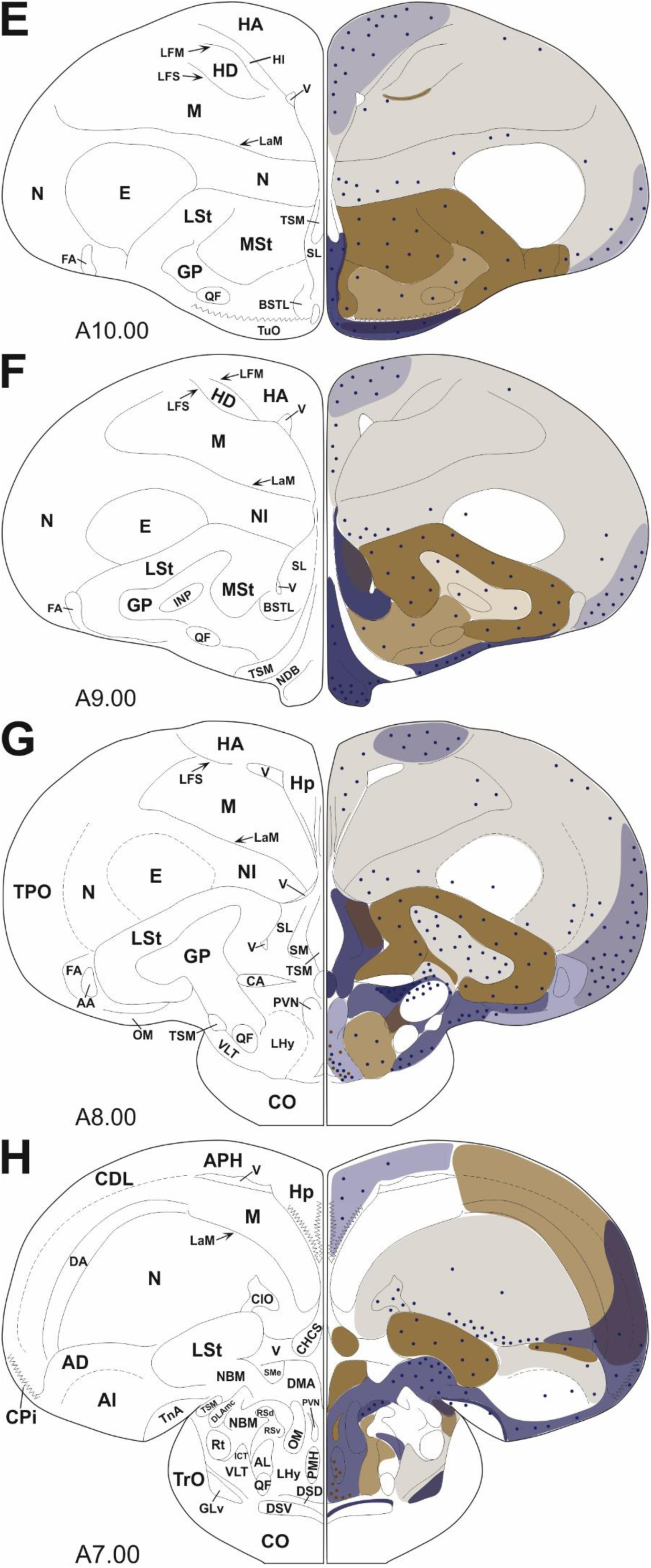
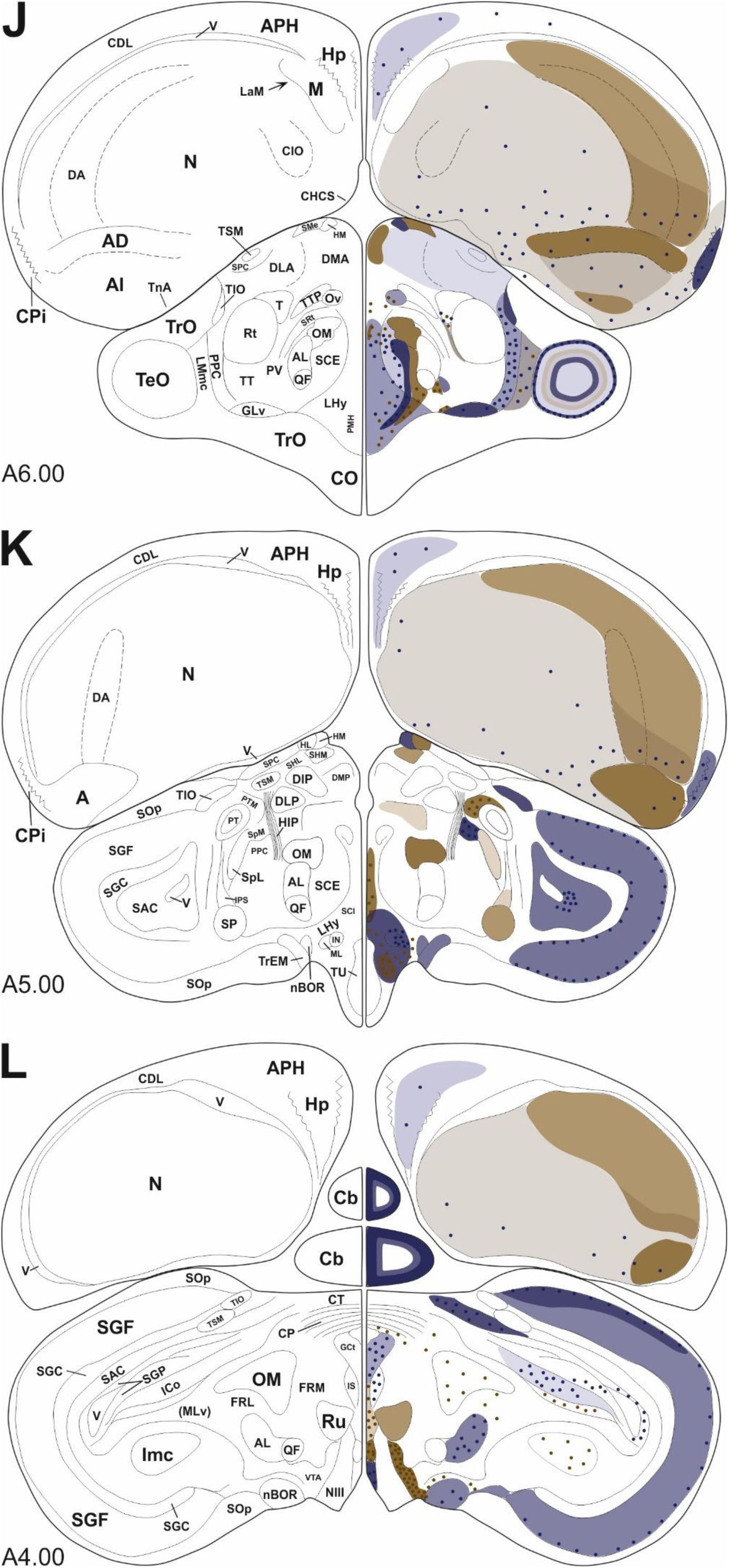
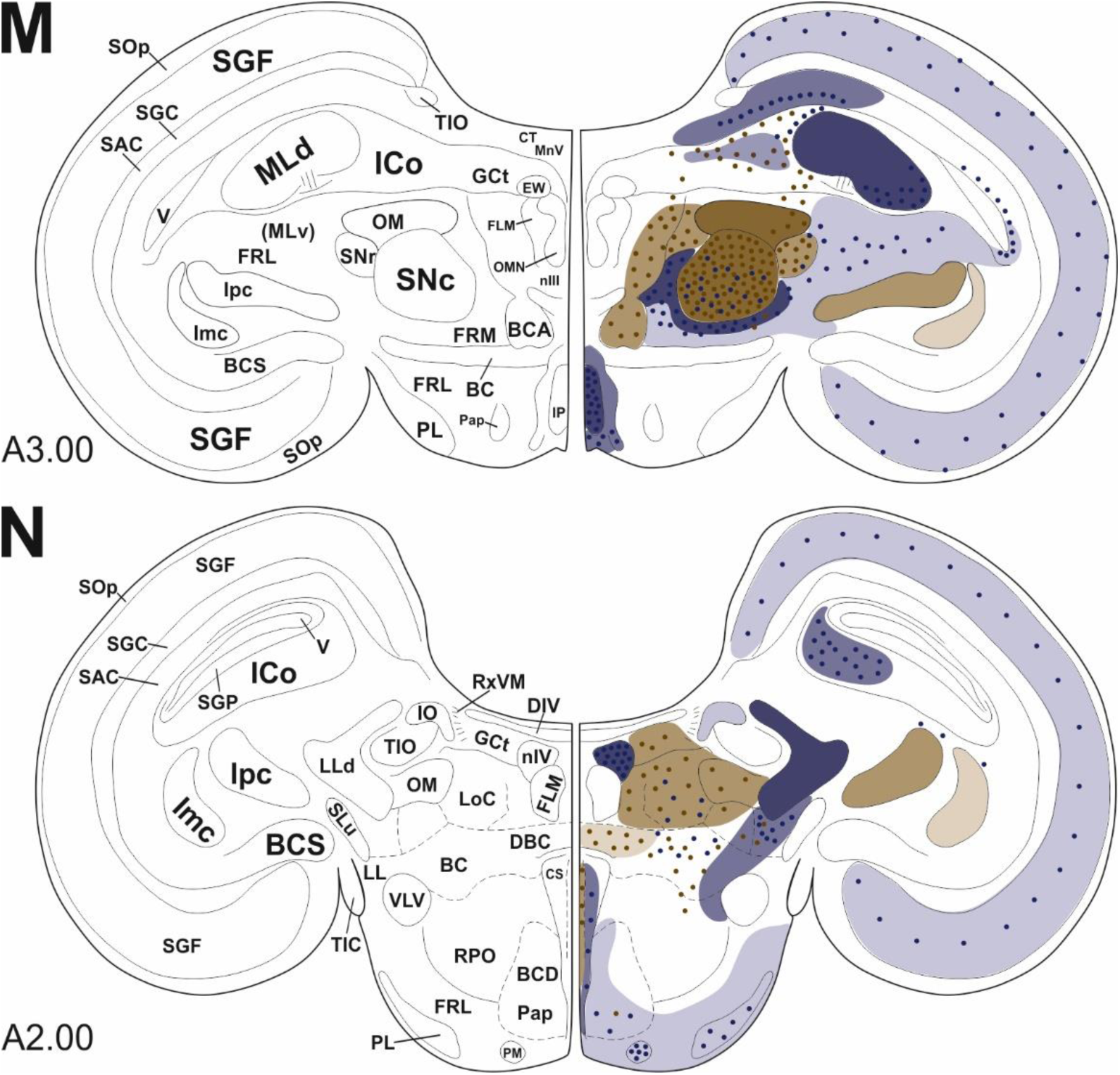
Schematic distributions of NADPH-d activity and TH immunoreactivity. For each slice, the left side displays the specific area denotations and the right side shows NADPH-d and TH labeling. NADPH-d activity is represented in blue and TH immunoreactivity in brown. Dots indicate labeled neurons and shaded areas indicate labeled fibers and neuropil. The number of dots does not represent the true but the relative number of labeled neurons. The intensity of a shaded area represents the density of fibers and neuropil with darker colors reflecting a higher density. For abbreviations, see Karten and Hodos (1967), Reiner et al. (2004), and list.

### Telencephalon

The olfactory bulb showed intense NADPH-d neuropil staining for the external plexiform layer (EPL) and mitral cell layer (MCL), and very intense staining for the internal granular layer (IGL) while the glomerular layer (GL) was spared (Figure 2 A and Figure 3 A, B). There were clearly no NADPH-d-positive neurons in the EPL or MCL. However, detection of labeled neurons in the IGL was not possible due to the very intense neuropil staining. Few weakly TH-positive neurons were found at the border between GL and EPL (Figure 3 B), and few weakly TH-positive fibers were found throughout all layers of the olfactory bulb. Moderate NADPH-d neuropil staining as well as both weakly and intensely stained neurons were observed in the prepiriform cortex (CPP; Figure 2 A and Figure 3 C). Few weakly TH-positive fibers and fiber baskets were also present here.

**Figure 3:**
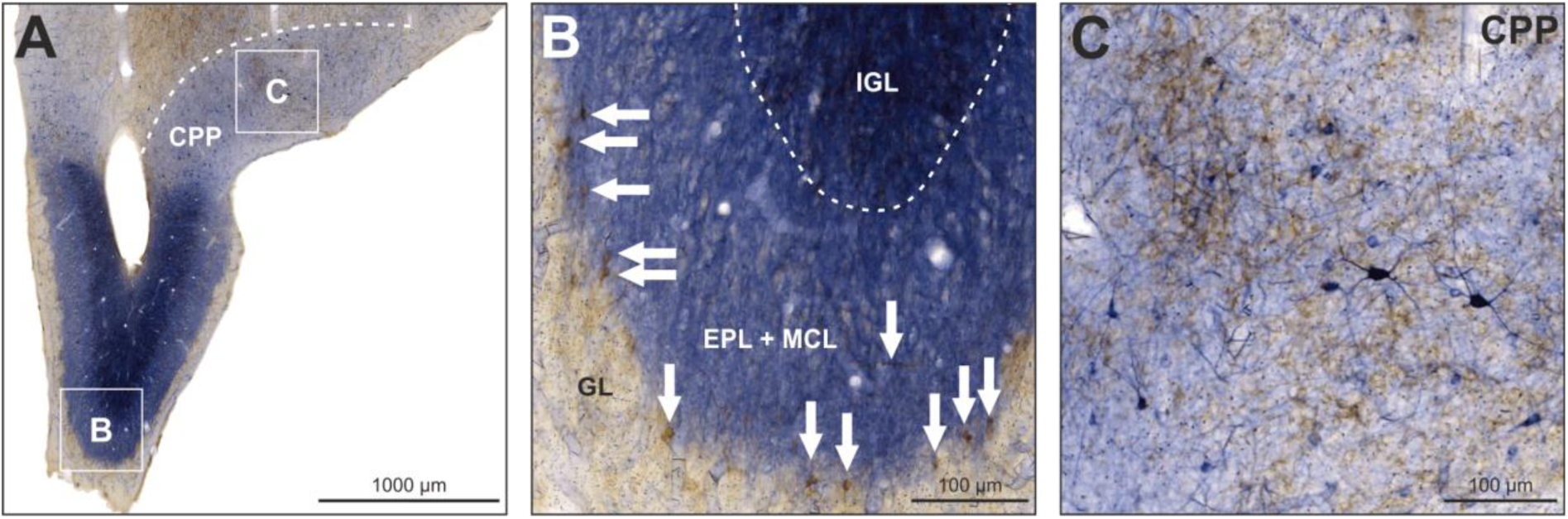
NADPH-d and TH in the olfactory bulb. NADPH-d is visualized in blue, TH in brown (upper panel), and NOS is visualized in black (lower panel). **(A)** The OB displayed intense NADPH-d activity in neuropil. **(B)** Enlargement of box B depicted in (A). The white arrows along the border between GL and EPL indicate TH-positive neurons and the arrow in EPL indicates a TH-positive fiber. **(C)** Enlargement of box C depicted in (A). Within CPP, neurons and neuropil positive for NADPH-d were detected as well as fibers and fiber baskets positive for TH.

Within the hyperpallium, especially the hyperpallium apicale (HA) displayed moderate NADPH-d activity in neuropil and a moderate number of labeled neurons, with the majority located in the medial and dorsal portions of HA (Figure 2 A-G and Figure 4 A, B). Among labeled neurons, there were bigger and intensely labeled ones with extensive axons and dendrites as well as smaller and less intensely labeled ones with only weakly labeled axons and dendrites if any. Very rarely, labeled neurons were also detected in the hyperpallium intercalatum (HI) and hyperpallium densocellulare (HD). The whole hyperpallium was weakly but evenly innervated by TH-positive fibers and baskets. Within the mesopallium, no noteworthy NADPH-d activity was detected except for some rare solitary neurons (Figure 2 A-J and Figure 4 A, C). In contrast to the hyperpallium, the mesopallium was characterized by a high density of TH-positive fibers and baskets (Figure 4 A, B). The lamina frontalis superior (LFS) constitutes the border between hyperpallium and mesopallium and displayed a very high density of TH-positive fibers.

**Figure 4:**
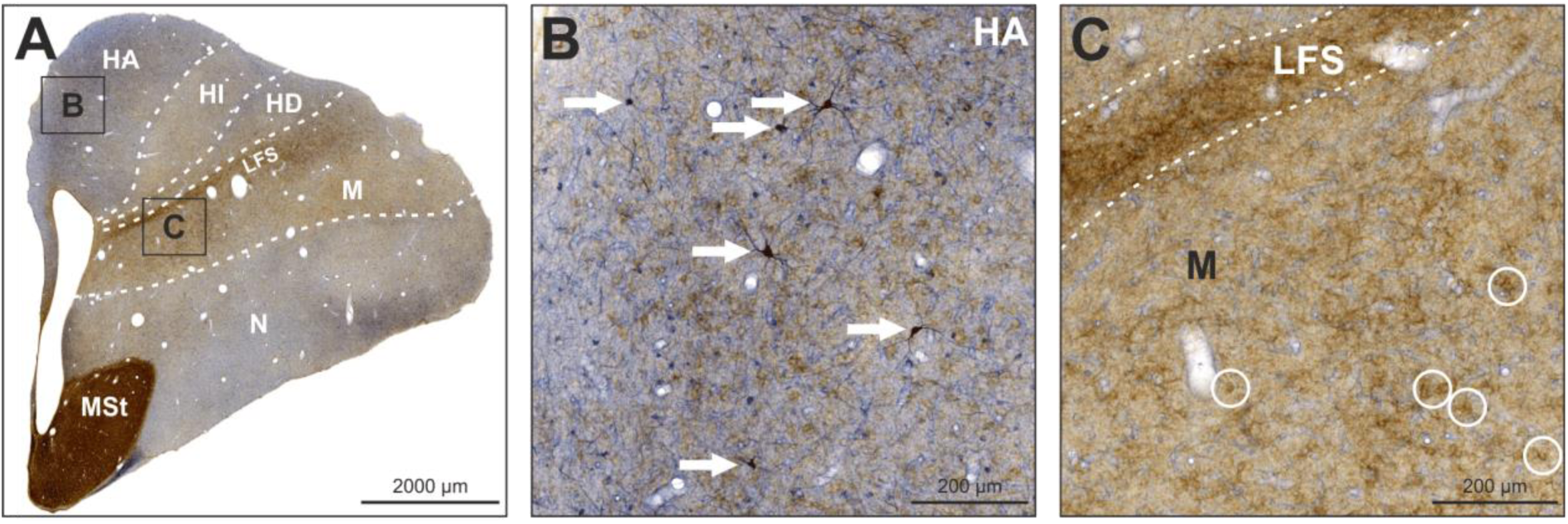
NADPH-d and TH labeling in the hyperpallium and mesopallium. NADPH-d is visualized in blue, TH in brown (upper panel), and nNOS is visualized in black (lower panel). **(A)** Frontal section at A13.00 stained for NADPH-d and TH. **(B)** Enlargement of box B depicted in (A). HA displayed moderate NADPH-d activity in both neuropil and neurons. White arrows indicate examples of intensely labeled neurons with axons and dendrites. Fibers and fiber baskets positive for TH are also visible. **(C)** Enlargement of box C depicted in (A). The mesopallium, especially the LFS, displayed a dense innervation by fibers and fiber baskets positive for TH. White circles indicate examples of fiber baskets. No NADPH-d activity is visible.

Within the nidopallium, labeled neurons were detected throughout its entire antero-posterior extent (Figure 2 A-L). In particular, neurons were predominantly located in the nidopallium intermedium medialis (NIM; Figure 5 B), nidopallium intermedium laterale (NIL; Figure 5 C), ventral nidopallium caudocentrale (NCC; Figure 5 E), piriform cortex (CPi), and temporo-parieto-occipital area (TPO; Figure 5 F). A substantial number of NADPH-d-positive neurons was also detected in the ventrolateral nidopallium dorsally adjacent to the arcopallium. Moderate to intense neuropil staining was observed for NIL, TPO, and CPi. Most of the nidopallium displayed a low density of TH-positive fibers and fiber baskets except for the nidopallium caudolaterale (NCL) which displayed a very high density of TH-positive baskets.

**Figure 5:**
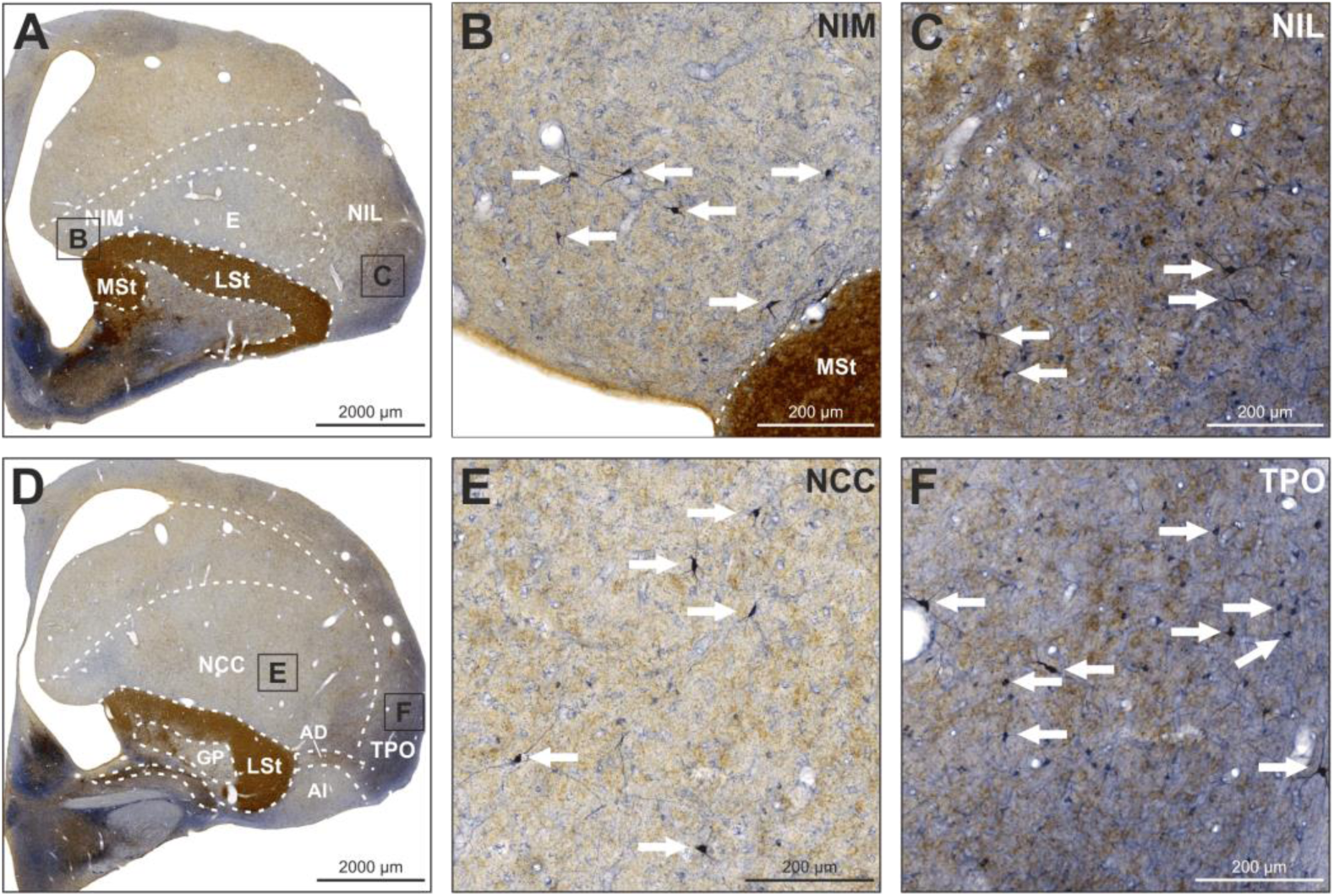
NADPH-d and TH, and nNOS labeling in the nidopallium. NADPH-d is visualized in blue, TH in brown (upper two panels), and nNOS is visualized in black (lower two panels). **(A)** Frontal section at A9.00 stained for NADPH-d and TH. **(B)** Enlargement of box B depicted in (A). NIM displayed weakly TH-positive fibers. White arrows indicate examples of intensely NADPH-d-positive neurons. **(C)** Enlargement of box C depicted in (A). Within NIL, a moderate density of TH-positive and NADPH-d-positive fibers was observed. White arrows indicate examples of intensely NADPH-d-positive neurons. **(D)** Frontal section at A7.50 stained for NADPH-d and TH. **(E)** Enlargement of box E depicted in (D). The NCC displayed a weak but even innervation by TH-positive fibers. NADPH-d-positive fibers were also visible. White arrows indicate examples of intensely NADPH-d-positive neurons. **(F)** Enlargement of box F depicted in (D). TPO displayed fibers positive for NADPH-d and TH. White arrows indicate examples of intensely NADPH-d-positive neurons.

Within the arcopallium, we observed weak NADPH-d activity in neuropil in the dorsal arcopallium (AD), increasing towards its medial portion (Figure 6 A). We did not observe NADPH-d-positive neuropil in the intermediate arcopallium (AI; Figure 6 C). Labeled neurons were found across all subdivisions and were mostly small and weakly stained (Figures 2 H-L and Figure 6 C). Few intensely stained neurons were scattered. While AD was heavily innervated by TH-positive fibers, AI showed only weak to moderate TH immunoreactivity (Figure 2 H-L and Figure 6 A, C). The posterior arcopallium also showed TH immunoreactivity in ventral AI. The adjacent nucleus taeniae of the amygdala (TnA) and nucleus posterioris amygdalopallii (PoA) were in general negative for both TH and NADPH-d (Figure 6 A, B). In some animals we detected small neurons that were faintly positive for NADPH-d in PoA. In some animals, the border between TnA and AI was positive for either TH or NADPH-d or both, and in some animals, we also observed NADPH-d positive neurons in this border region.

**Figure 6:**
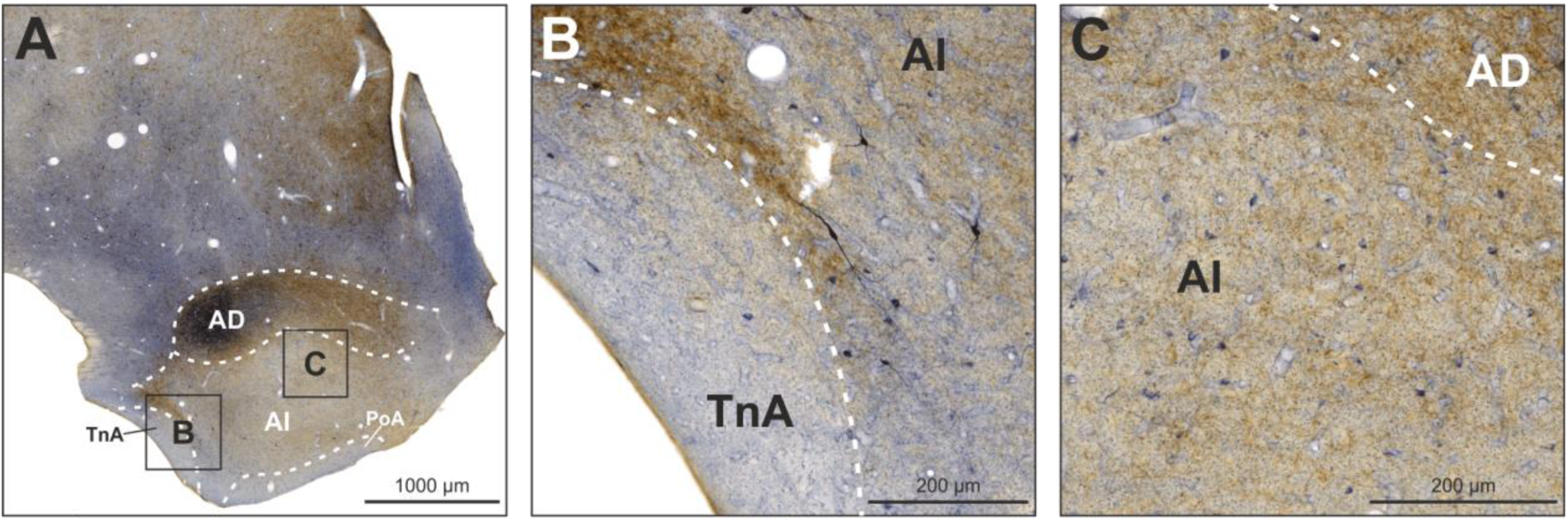
NADPH-d and TH, and nNOS labeling in the arcopallium and adjacent limbic structures. NADPH-d is visualized in blue, TH in brown (upper panel), and nNOS is visualized in black (lower panel). **(A)** Frontal section at A6.50 stained for NADPH-d and TH. **(B)** Enlargement of box B depicted in (A). TnA is mostly negative for both NADPH-d and TH while AI displays neurons intensely and weakly positive for NADPH-d. **(C)** Enlargement of box C depicted in (A). AI displayed moderate TH immunoreactivity and evenly distributed NADPH-d-positive neurons

The entopallium did not show NADPH-d activity in neuropil and labeled neurons were detected very rarely (Figure 2 D-G and Figure 7 A-B). TH-positive fibers and fiber baskets were very sparse but evenly distributed. The striatum was heavily innervated by TH positive fibres so that the NADPH-d staining was hard to assess. Therefore, we also stained some sections exclusively against NADPH-d to investigate neuropil and neurons in these regions. We found a moderate numbers of NADPH-h positive neurons in MSt, LSt and GP (Figure 7 C).

**Figure 7:**
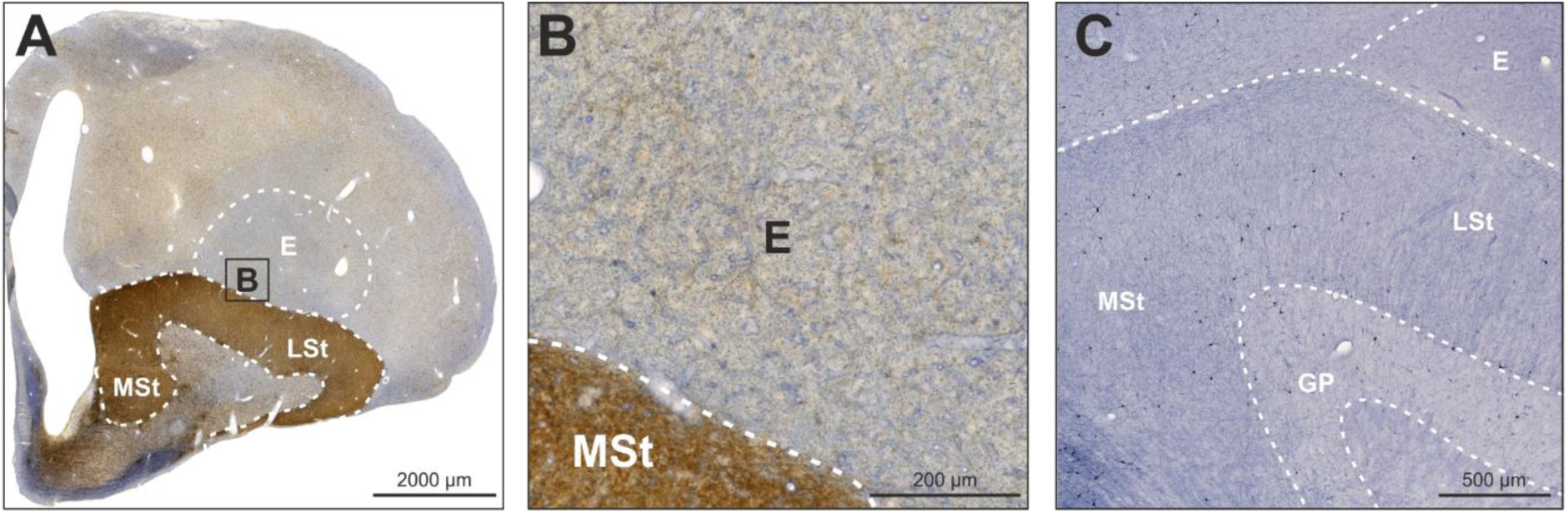
NADPH-d and TH abeling in the entopallium. NADPH-d is visualized in blue, TH in brown (upper panel), and nNOS is visualized in black (lower panel). **(A)** Frontal section at A9.50 stained for NADPH-d and TH. **(B)** Enlargement of box B depicted in (A). The entopallium showed weak immunoreactivity for TH but was negative for NADPH-d. (C) The striatum showed moderate immunoreactivity for NADPH-d.

Within the hippocampus, neuropil was moderately positive for NADPH-d, particularly in the triangular region (Tr), and to a lesser extent also in the dorsal and ventral portions of the dorsomedial area (DMd, DMv; Figure 2 H-L and Figure 8 A, B). NADPH-d-positive neurons were found in all hippocampal subregions as well as in the area corticoidea dorsolateralis (CDL; Figure 2 H-L and Figure 8 A). Moreover, we detected a cluster of intensely stained TH-positive fibers that was located in DMd and extended into the dorsal portion of DMv (DMvd) in some animals (Figure 8 C). In some animals this cluster also contained few TH-positive neurons.

**Figure 8:**
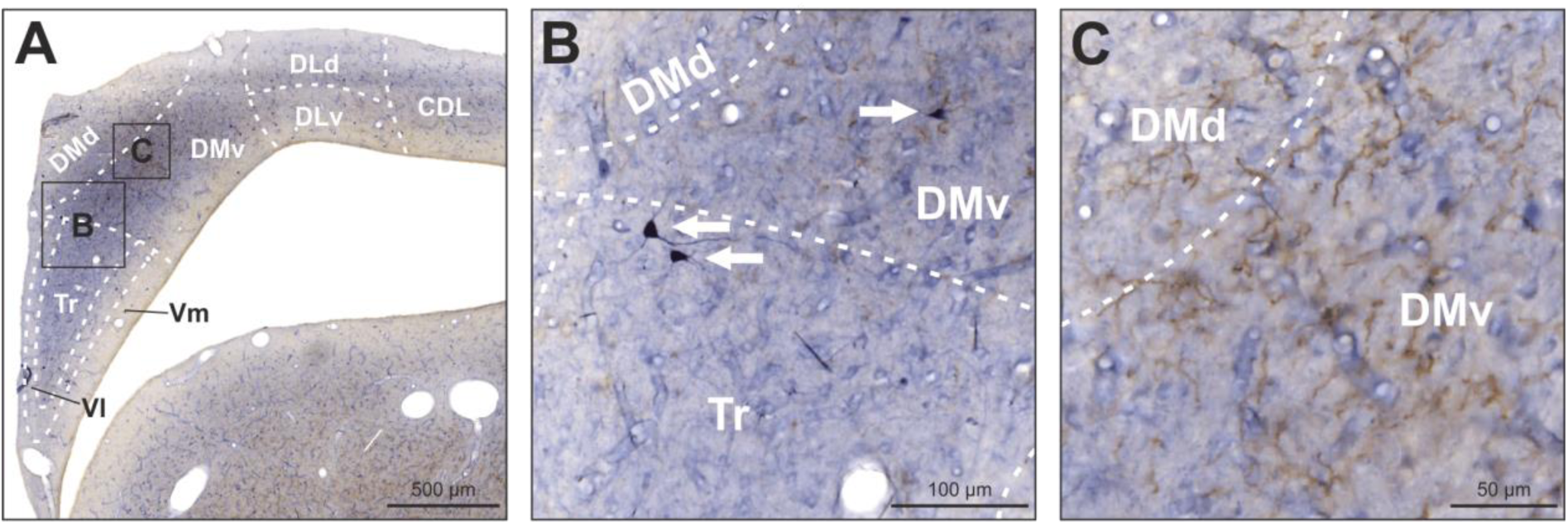
NADPH-d and TH labeling in the hippocampus. NADPH-d is visualized in blue, TH in brown (upper panel), and nNOS is visualized in black (lower panel). **(A)** Frontal section at A5.00 stained for NADPH-d and TH. **(B)** Enlargement of box B depicted in (A). NADPH-d-positive neuropil was observed in Tr, DMd, and DMv. White arrows indicate examples of intensely NADPH-d-positive neurons. **(C)** Enlargement of box C depicted in (A). In DMd and DMv, intensely TH-positive fibers were detected.

Within the basal ganglia, both the lateral (LSt) and medial striatum (MSt) contained a moderate number of NADPH-d-positive neurons with the majority located at the periphery towards the nidopallium, entopallium, or globus pallidus (GP; Figure 2 B-H, Figure 7 C and Figure 9 C, H). The GP also contained a moderate number of intensely NADPH-d-positive neurons (Figure 2 F, G and Figure 9 D, E). In some animals, these were primarily located in the dorsal portion along the border between GP and LSt. GP fibers were positive for both NADPH-d and TH. The lateral forebrain bundle (LFB) displayed weakly NADPH-d-positive fibers in the dorsolateral portion and intensely positive fibers in the ventromedial portion. We also observed a cluster of intensely labeled neurons along the medial border of the LFB (Figure 9 D, G). Moreover, we observed intense neuropil staining in the adjacent nucleus septalis medialis (SM), nucleus preopticus anterior (POA), and tuberculum olfactorium (TuO). In the POA and TuO, we also detected labeled neurons. The nucleus of the diagonal band (NDB) displayed fibers positive for both NADPH-d and TH (Figure 9 J). The nucleus accumbens (Ac) was heavily innervated by TH-positive fibers and also displayed few NADPH-d-positive neurons. In the dorsal portion of the lateral bed nucleus of the stria terminalis (BSTL), we observed a small band of NADPH-d-positive neurons and neuropil (Figure 9 D, F). Furthermore, TH-positive fibers were found in the TuO, POA, ventral LFB, ventral pallidum (VP), nucleus septalis lateralis (SL), nucleus commissuralis septi (CoS), and tractus cortico-habenularis et cortico septalis (CHCS). Because of the heavy TH immunoreactivity, it was very difficult to detect any potential NADPH-d activity in Ac or SL

**Figure 9:**
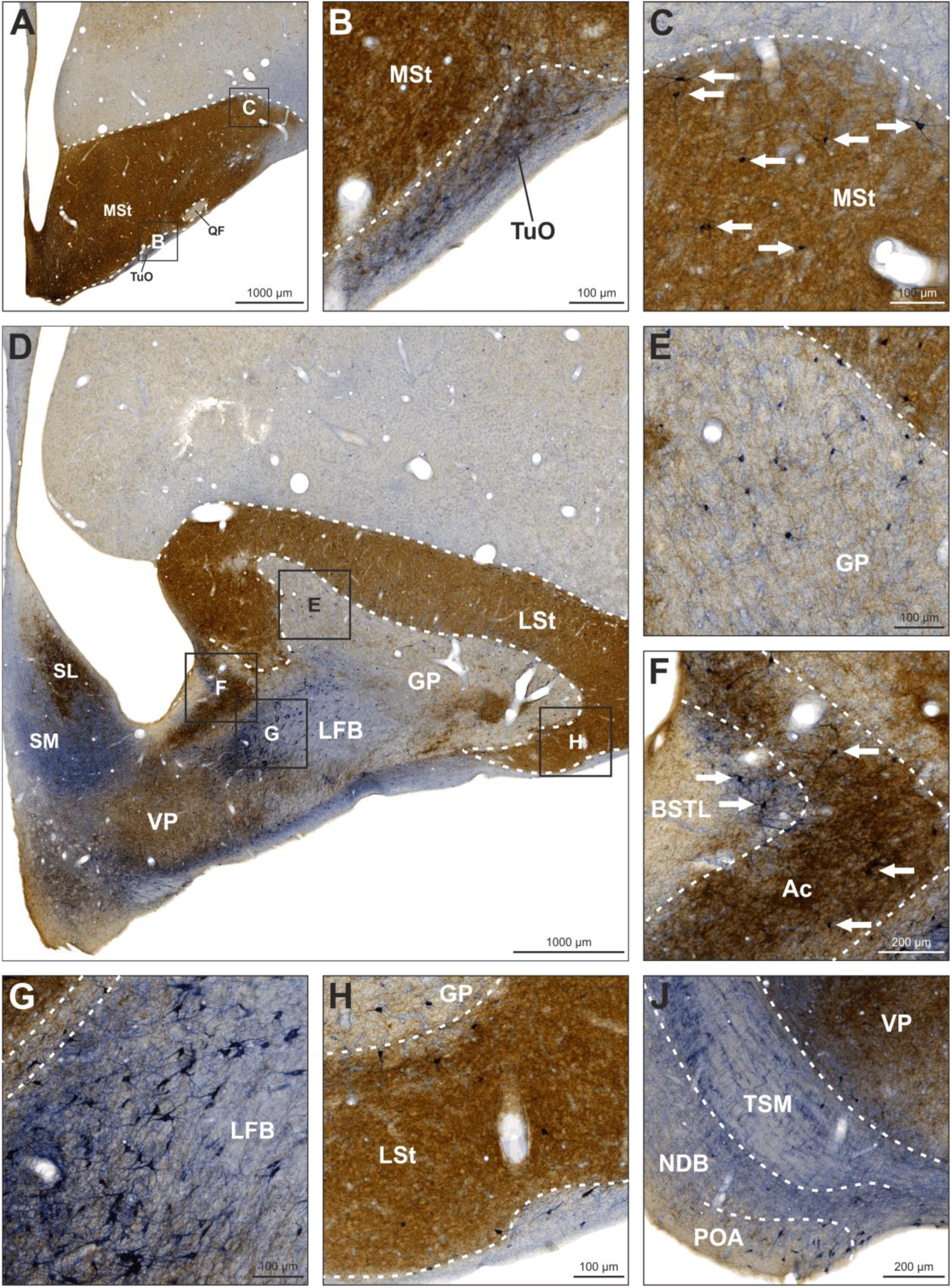
NADPH-d and TH labeling in the basal ganglia and adjacent structures. NADPH-d is visualized in blue and TH in brown. **(A)** Frontal section at A10.50. **(B)** Enlargement of box B depicted in (A). Fibers in the TuO were positive for both NADPH-d and TH. Neurons weakly positive for NADPH-d were also detected. **(C)** Enlargement of box C depicted in (A). The MSt was heavily innervated by TH-positive fibers. White arrows indicate neurons intensely positive for NADPH-d. **(D)** Frontal section at A8.00. **(E)** Enlargement of box E depicted in (D). The GP contained a moderate number of NADPH-d-positive neurons, and fibers positive for both NADPH-d and TH. **(F)** Enlargement of box F depicted in (D). The Ac was heavily innervated by TH-positive fibers and exhibited few NADPH-d-positive neurons as indicated by white arrows. The dorsolateral portion of the adjacent BSTL displayed a thin band of NADPH-d-positive fibers and neurons as indicated by white arrows. **(G)** Enlargement of box G depicted in (D). The dorsomedial portion of the LFB showed intensely NADPH-d-positive neurons and fibers. **(H)** Enlargement of box H depicted in (D). The LSt was heavily innervated by TH-positive fibers and contained a moderate number of NADPH-d-positive neurons. **(J)** Frontal section at A9.00. The VP was heavily innervated by TH-positive fibers. The TSM showed NADPH-d activity in fibers running along the tract as well as in fibers crossing orthogonally. NDB showed fibers positive for NADPH-d and TH, and NADPH-d-positive neurons in the ventrolateral portion. POA contained a high number of NADPH-d-positive neurons and fibers positive for NADPH-d and TH. For abbreviations, see list.

#### Midbrain

Within the midbrain, numerous nuclei were found positive for either NADPH-d or TH or both (Figure 2 L-N). The nucleus of the basal optic root (nBOR) displayed weak to moderate NADPH-d activity in neuropil, and only very few neurons (Figure 10 A). Neurons and fibers in the ventral tegmental area (VTA) were intensely positive for TH (Figure 10 A, B, D). Since the TH staining was so strong in VTA it was hard to assess the NADPH-d labeling. Therefore, we selectively stained some sections against NADPH-d- and found a large amount of NADPH-d positive neurons within VTA (Figure 10 E). Medial to the nervus occulomotorius (NIII) and dorsal to the nucleus interpeduncularis (IP), we detected more TH-positive fibers. IP contained intensely NADPH-d-positive neuropil and some neurons (Figure 10 B, F). The nucleus ruber (Ru) contained TH-positive fibers but was negative for NADPH-d (Figure 10 D). The nucleus interstitialis (IS) was easily distinguishable from its surroundings based on weak NADPH-d staining in neuropil and a moderate number of labeled neurons (Figure 10 C). Occasionally, we detected TH-positive neurons and fibers within the medial portion of IS, but it is likely that these belong to adjacent D. The substantia grisea centralis (GCt) mainly contained TH-positive neurons and fibers, but we also observed weakly NADPH-d-positive neuropil and few neurons (Figure 10 C, F, G). QF exhibited weakly TH-positive fibers (Figure 10 D). Intensely NADPH-d-positive neurons were found in AL extending into the formation reticularis lateralis mesencephalic (FRL) and OM, which predominantly contained TH-positive fibers and neurons (Figure 10 D). The formation reticularis medialis mesencephalic (FRM) mainly contained weakly NADPH-d-positive neurons and weakly TH-positive fibers (Figure 10 C, D). The nucleus mesencephalicus lateralis, pars dorsalis (MLd) showed an interesting pattern of NADPH-d activity. In none of the animals, the whole nucleus showed reactivity. In the anterior portion of MLd, we observed two discrete clusters of NADPH-d activity. The ventral cluster was usually bigger with more intense staining in neuropil and neurons, whereas the smaller dorsal cluster only showed activity in neuropil. In the posterior portion of MLd, activity was mainly concentrated at the ventrolateral tip and along the dorsal and ventral borders of the nucleus (Figure 10 G). The nucleus intercollicularis (ICo) showed moderate NADPH-d activity in neuropil and neurons, especially in its posterior portion. In its intermediate portion, where ICo is located between GCt and OM, we also detected many TH-positive neurons and fibers in its ventral half (Figure 10 F, G). The nucleus papilloformis (Pap) contained many moderately NADPH-d-positive neurons and fibers and also TH-positive fibers and few neurons. The nucleus mesencephalicus profundus, pars ventralis (MPv) contained TH-positive fibers and big intensely NADPH-d-positive neurons (Figure 10 B). As expected and similar to the VTA, neurons and fibers in the substantia nigra (SN) were heavily positive for TH, also encompassing adjacent areas (Figure 10 F, G). We also observed NADPH-d activity in neurons and fibers throughout the SN. The nucleus centralis superior (CS) was intensely positive for NADPH-d in neuropil and some neurons (Figure 10 H). The nucleus linearis caudalis (LC), which is located medially to the CS, displayed a thin band of TH-positive fibers and neurons (Figure 10 H). Alongside VTA and SN, the locus ceruleus (LoC) was the third major structure that contained intensely TH-positive neurons and fibers (Figure 10 J). Beneath this dense web, we also spotted NADPH-d-positive neurons. The brachium conjunctivum descendens (BCD) displayed a network of NADPH-d-positive fibers but no neurons. Furthermore, the nucleus lemnisci lateralis, pars dorsalis (LLd) showed very intense NADPH-d activity in neuropil (Figure 10 J). The nucleus isthmo-opticus (IO) showed weak to moderate NADPH-d activity in neuropil (Figure 10 J). The nucleus nervi trochlearis (nIV) displayed a dense network of NADPH-d-positive fibers and neuropil (Figure 10 J). The nucleus also contained intensely stained neurons and TH-positive fibers. In some, but not all birds, we observed NADPH-d activity in neuropil the in nucleus semilunaris (SLu, Figure 10 J). The nucleus ventmtis lemnisci lateralis (VLV) was negative for both NADPH-d and TH. The nucleus pontis lateralis (PL) and nucleus pontis medials (PM) in the ventral midbrain both showed moderate NADPH-d activity in neuropil. In PM, a high number of NADPH-d-positive neurons clustered, whereas in PL, a moderate number of neurons was distributed throughout the nucleus (Figure 10 K). A network of NADPH-d-positive neurons and fibers was widely distributed across the brachium conjunctivum descends et tractus tectospinalis (BCTS), nucleus reticularis pontis oralis (RPO), nucleus reticularis pontis caudalis, pars gigantocellularis (RPgc), and corpus trapezoideum (CTz, Figure 10 L).

**Figure 10:**
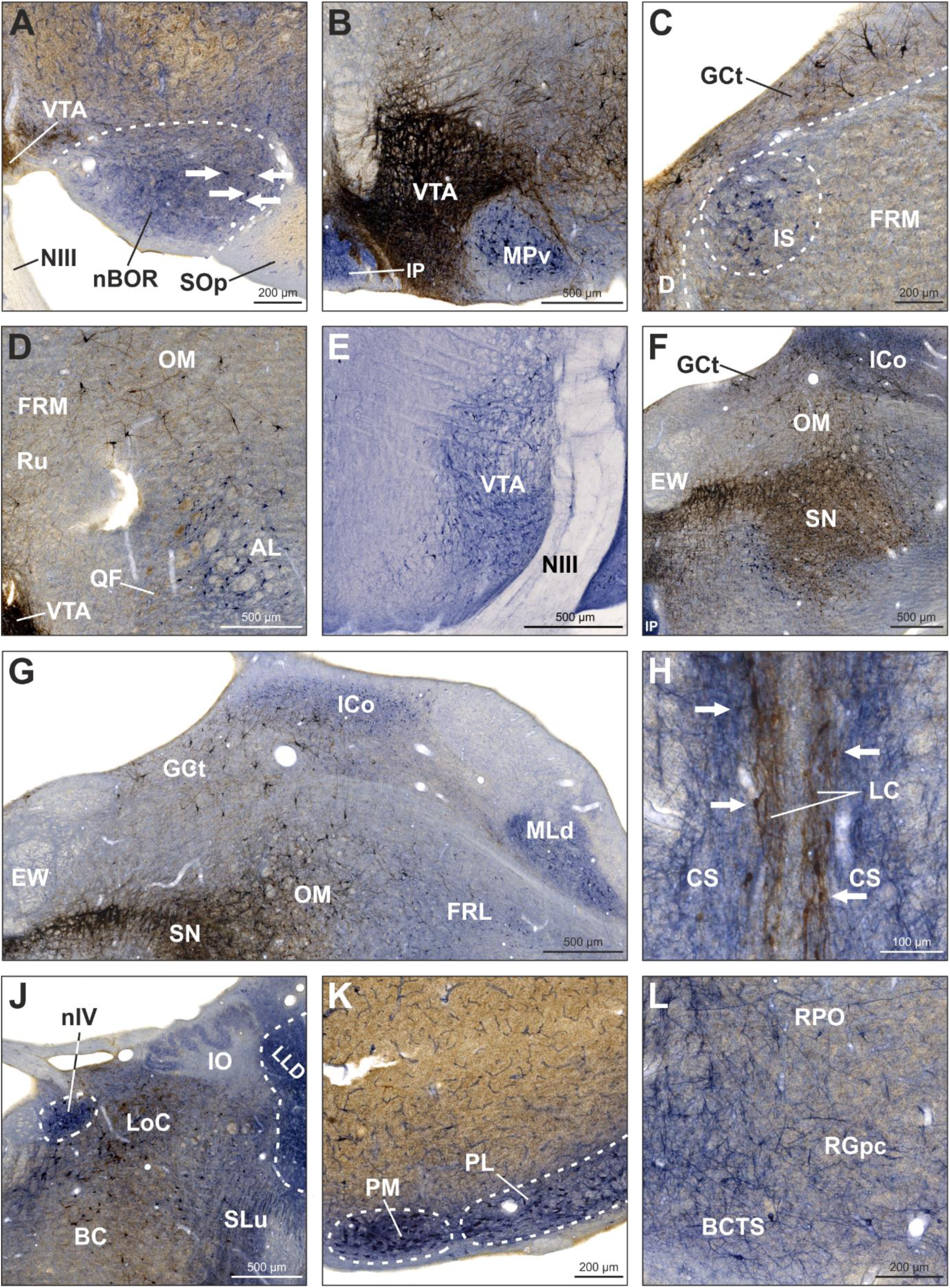
NADPH-d and TH labeling in the midbrain. NADPH-d is visualized in blue and TH in brown. All pictures depict right hemispheres except for (H) which depicts a section along the centerline of the slice including both hemispheres. **(A)** Frontal section at A4.00. Neuropil in the nBOR was moderately positive for NADPH-d. White arrows indicate labeled neurons. **(B)** Frontal section at A3.50. VTA contained a dense network of TH-positive neurons and fibers. NADPH-d activity in the VTA was also observed but difficult to distinguish due to the strong TH staining intensity. Neuropil and neurons in IP and MPv were NADPH-d-positive. **(C)** Frontal section at A4.00. IS contained neuropil and neurons positive for NADPH-d. D and GCt showed TH-positive neurons and fibers. FRM contained weakly TH-positive fibers and weakly NADPH-d-positive neurons. **(D)** Frontal section at A4.00. VTA contained a dense network of TH-positive neurons and fibers. QF showed weakly TH-positive fibers. Intensely NADPH-d-positive neurons were distributed across AL and adjacent areas. Ru contained TH-positive fibers. Intensely TH-positive neurons were distributed across OM. **(E)** Frontal section showing NADPH-d positive neurons within VTA. **(F)** Frontal section at A3.00. SN was intensely TH-positive in neurons and fibers. NADPH-d-positive neurons were detected in the ventral SN and extended beyond its medial border. OM displayed TH-positive neurons and fibers extending into GCt and ventral ICo. ICo also showed NADPH-d activity in neurons and neuropil. **(G)** Frontal section at A3.00. Dorsal ICo showed NADPH-d activity in neuropil and neurons as well as TH-positive fibers. Ventral ICo predominantly contained TH-positive neurons and fibers. MLd showed NADPH-d activity in neuropil, neurons and fibers in its ventrolateral tip. **(H)** Frontal section at A2.25. LC contained TH-positive fibers and neurons. White arrows indicate examples of labeled neurons. CS contained NADPH-d-positive fibers. In other cases, we also observed NADPH-d-positive neurons. **(J)** Frontal section at A2.00. LoC displayed TH immunoreactivity in neuropil, neurons, and fibers, and NADPH-d activity in neurons and fibers. nIV was intensely NADPH-d-positive. TH-positive neurons and fibers were distributed across BC. Neuropil in LLd was intensely NADPH-d-positive. SLu showed NADPH-d activity in neuropil and neurons and few TH-positive fibers. **(K)** Frontal section at A1.75. PM and PL contained intensely NADPH-d-positive neuropil and a high number of labeled neurons. **(L)** Frontal section at A1.50. A network of NADPH-d-positive fibers and few neurons was distributed across RPO, RGpc, and BCTS. For abbreviations, see list.

### Colocalization of nNOS and TH

Thus far, we investigated the distributions of NADPH-d and TH separately which did not allow for a precise detection of colocalizations. However, in order to assess the neuroanatomical framework of the memory flexibility theory that has been established in fruit flies we needed a combined investigation of TH, NOS and sGC. We therefore conducted fluorescence stainings with tissue of six animals to determine where TH and NOS overlap within the midbrain and whether TH and NOS positive fibres target sGC-positive cells in the forebrain.

All of the major TH-positive midbrain structures, the LoC, VTA, and the SN showed comparable colocalization of TH and nNOS (Figure 11 A-J). In addition, we found that many TH-positive neurons in the GCt also expressed nNOS (Figure 11 K-M).

**Figure 11:**
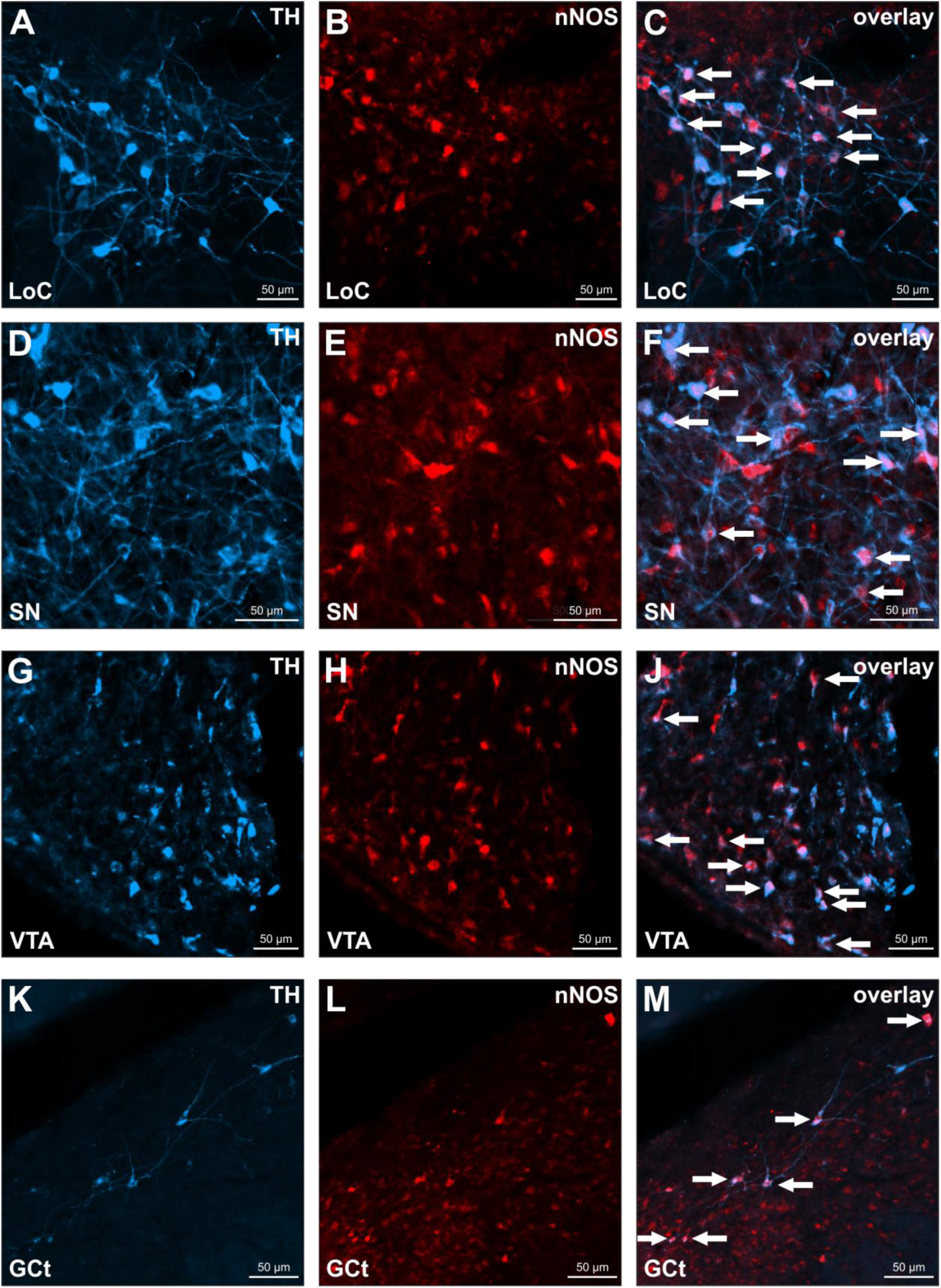
Colocalization of nNOS and TH. Each row is dedicated to one TH-positive midbrain structure (from top to bottom LoC, SN, VTA, GCt). Left column pictures depict TH-positive neurons in blue, middle column pictures depict nNOS-positive neurons in red at the same site, right column pictures depict overlays. White arrows indicate exemplary neurons positive for both TH and nNOS. For abbreviations, see list.

Furthermore, we investigated the anatomical relationships among TH-positive, NADPH-d positive afferent fibers and sGC-positive cells in the pigeon nidopallium caudolaterale (NCL), which is one of the primary target of dopaminergic input in the avian telencephalon and commands executive functions (Figure 2). Therefore. we conducted a triple staining protocol, in which sGC-positive cells were permanently stained brown using standard DAB staining (Figure 12 A,B), NOS-positive fibers were labeled blue through NADPH staining (Figure 12 A,B), and TH-positive fibers were visualized in red using fluorescence staining (Figure 12 C). Our results reveal that both NOS- and TH-positive fibers appear to encircle sGC-positive cells, indicating a close spatial association (Figure 12 D).

**Figure 12:**
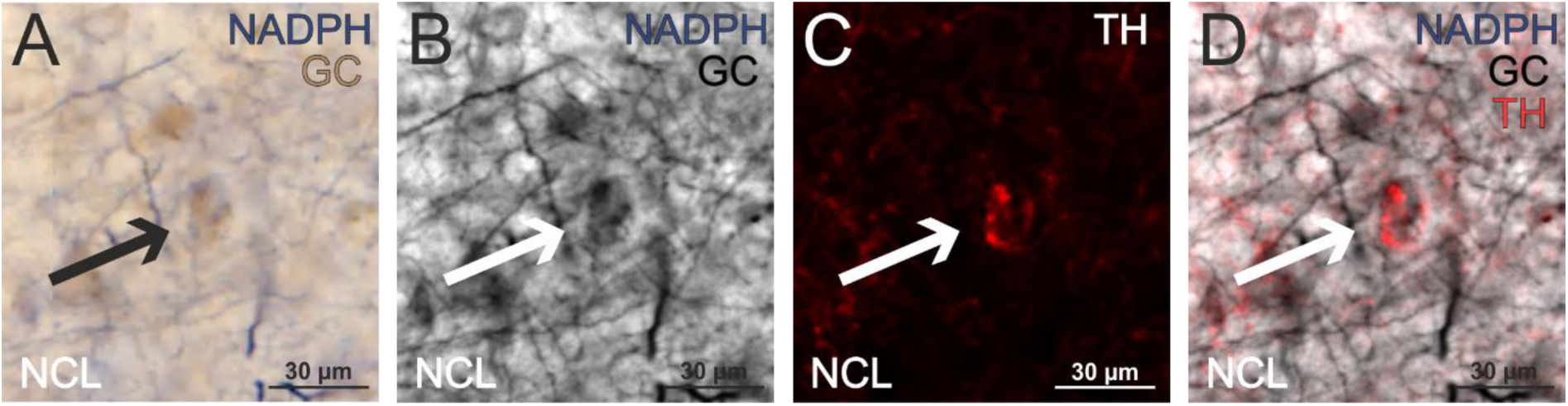
Spatial Association of TH, NOS, and sGC-Positive Cells in the Pigeon NCL using Triple Staining. **(A)** displays NOS-positive fibers in blue and sGC-positive cells in brown. **(B)** presents the same region in grayscale, showing NOS and sGC staining without color. **(C)** shows a fluorescence image where TH-positive fibers are depicted in red, highlighting their distribution in the NCL. **(D)** overlays the red TH fluorescence from panel (c) on the grayscale image from panel (b), demonstrating how both NOS and TH fibers appear to encircle the sGC-positive cells.

## Discussion

This study provides new insights into the distribution of NO-synthesizing neurons in the pigeon brain and their potential interactions with dopaminergic pathways, particularly in relation to memory flexibility. By mapping NADPH-d, nNOS, sGC, and TH, we identified a neuroanatomical framework supporting NO-dopamine interactions. These findings support the view that NO might play a role in modulating cognitive processes such as memory adaptation, complementing dopamine’s established functions in reinforcement learning. In particular, the structural convergence of NO and dopamine markers in the nidopallium caudolaterale (NCL) suggests a potential mechanism for regulating the balance between memory consolidation and forgetting, a feature reminiscent of NO’s function in *Drosophila melanogaster* (Aso et al., 2019).

### NO and Memory Flexibility

Memory is a dynamic process, requiring both stabilization and modification of existing traces. NO has been implicated as a key modulator of memory flexibility in Drosophila, where it co-releases with dopamine in select neurons, promoting gradual memory decay while allowing new learning to take place (Aso et al., 2019; Green & Lin, 2020). Experimental work in Drosophila has demonstrated that NO-producing dopaminergic neurons in the mushroom body regulate memory stability by weakening associations over time. Aso et al. (2019) found that genetic inhibition of NO synthase in Drosophila led to an over-stabilization of memories, preventing their expected decay, while NO signaling facilitated memory updating and reduced retention duration (Figure 13A). This indicates that NO functions as a regulatory signal, ensuring that memories do not persist indefinitely but instead remain adaptable to environmental changes. In our study, we show that within the NCL, both dopaminergic projections and NADPH-d-positive fibers converge on the same sGC-positive neurons (Figures 12, 13B). Building on findings from Drosophila, this structural arrangement suggests that NO in pigeons may also contribute to the modification of previously learned associations, which would be particularly relevant in extinction learning paradigms that support adaptive behavior (Packheiser et al., 2021).

**Figure 13.**
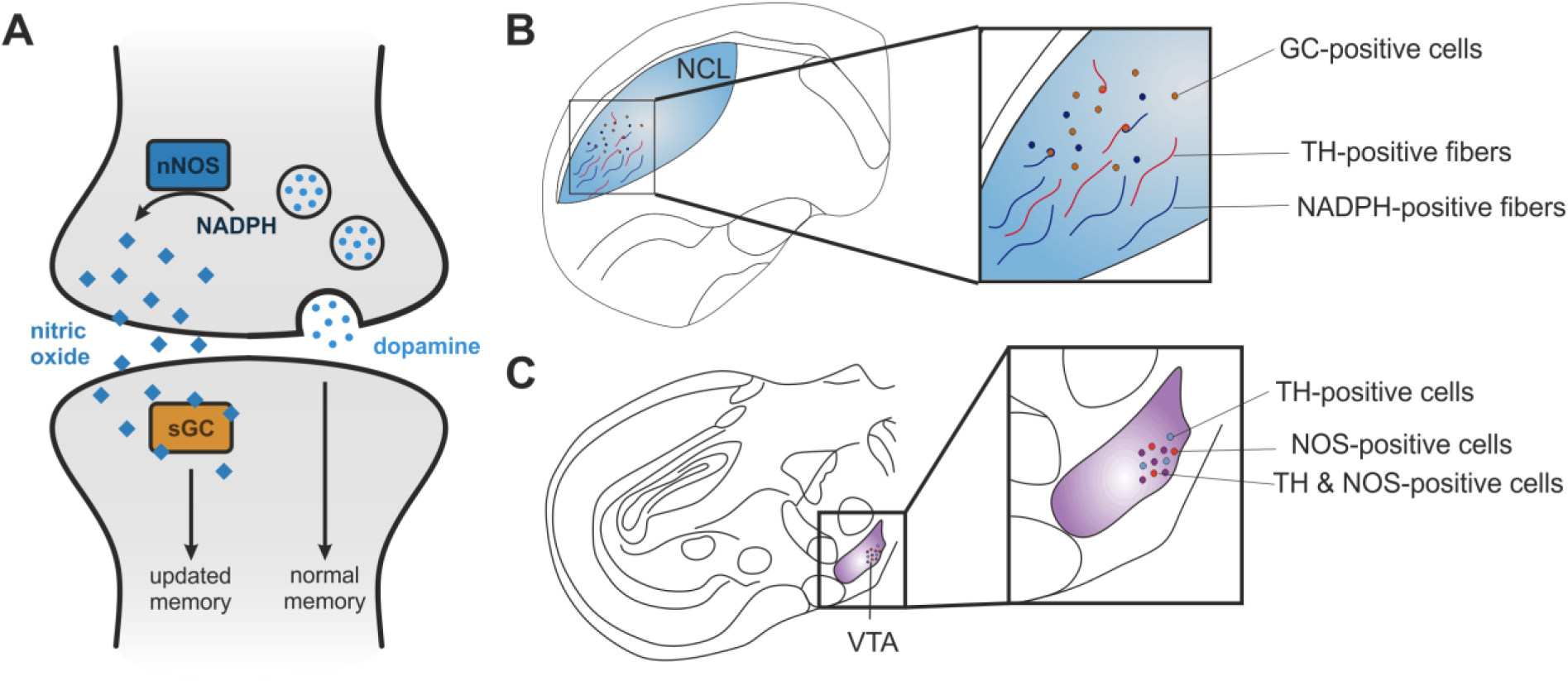
Summary of the memory flexibility hypothesis and our main findings. **(A)** in *Drosophila melanogaster* a subset of dopaminergic neurons releases both dopamine and nitric oxide (NO), enabling the fly to both reinforce and gradually “forget” memories associated with rewards or punishments (Aso et al., 2019; Green & Lin, 2020). In this system, dopamine strengthens memory traces tied to meaningful experiences, while NO serves as a complementary, slowly acting signal that weakens these memories over time. **(B)** In pigeons, a similar neuroanatomical framework can be found as some TH and NADPH-d positive fibres terminate on sGC positive cells within the NCL. **(C)** In pigeons, a similar neuroanatomical framework can be found as some neurons within the VTA express both NOS and TH.

The interaction between NO and dopamine is further supported by our observation that nNOS-positive neurons are present in the VTA, LoC, and SN, with no significant difference in colocalization levels with TH between these regions (Figure 13C). This implies that NO may influence multiple dopaminergic pathways, aligning with previous studies in mammals where NO affects synaptic plasticity and dopamine release in various regions, including the prefrontal cortex (Bon & Garthwaite, 2003; Kiss et al., 1999; Pogun et al., 1994; Trabace et al., 2007; West et al., 2002). Overall, the importance of NO in regulating synaptic plasticity (Bon and Garthwaite 2003; Böhme et al. 1993), along with its presence in areas associated with decision-making and response adaptation (Kiss et al., 1999; Trabace et al., 2007; Zoubovsky et al., 2011) supports the idea that NO plays a fundamental role in modifying pre-existing behavioral strategies based on environmental demands.

Dopaminergic pathways are essential for working memory, reinforcement learning, cognitive flexibility, and decision-making (Karakuyu et al., 2007; Packheiser et al., 2021; Puig et al., 2014; Pusch et al., 2023; Rook et al., 2023). In pigeons, the NCL is a primary target for dopaminergic input, with TH-positive fibers coiling around non-dopaminergic neurons (Divac et al., 1985; Kitt & Brauth, 1986; Waldmann & Güntürkün, 1993; Wynne & Güntürkün, 1995). The convergence of NO-synthesizing and TH-positive fibers on sGC-expressing neurons in the NCL (Figure 13B) reinforces the idea that NO may modulate dopamine-driven cognitive functions in birds as well. Comparable mechanisms have been described in mammals, where NO mediated changes in prefrontal dopamine release affect working memory and learning (Trabace et al., 2007; Zoubovsky et al., 2011). Moreover, research in rodents has shown that NO acts as a neuromodulator of learning and memory processes, particularly those requiring adaptation to new contingencies. Evidence from both inhibitory avoidance and spatial learning tasks suggests that NO plays a role in memory updating, behavioral flexibility, and reinforcement-based learning mechanisms. Inhibitory avoidance paradigms have demonstrated that modulation of NO in the hippocampus affects both short- and long-term memory retention, indicating a role in regulating long-term memory stability (Harooni et al., 2009). Likewise, NO inhibition impairs performance during the acquisition phase of the radial arm maze—a task requiring animals to integrate spatial cues and to avoid revisiting previously explored locations—revealing a role in spatial working memory and adaptive strategy use (Yamada et al., 1995). Furthermore, NO inhibition has been shown to alter dopamine metabolism in the striatum, implying that NO interacts with dopamine to modulate reinforcement-based learning mechanisms. Together, these findings highlight NO as a conserved neuromodulator involved in regulating synaptic plasticity and shaping adaptive learning processes across species, with our results suggesting a similar role in the avian brain.

### NO in Spatial Memory and Aging

Beyond its role in memory flexibility, NO has also been implicated in spatial memory formation and navigation (Estall et al., 1993; Majlessi et al., 2008; Noda et al., 1997; Yamada et al., 1995; Zou et al., 1998). The presence of NO-synthesizing neurons in the pigeon hippocampus is consistent with the idea that NO contributes to spatial learning, as has been shown in rodents, where NO is essential for hippocampus-dependent navigation tasks (Majlessi et al., 2008; Zhang et al., 1998). As pigeons rely heavily on spatial memory for homing and foraging (Herold et al., 2015; Krebs et al., 1996), NO may serve as a regulatory factor ensuring that spatial representations remain adaptable over time. In light of these findings, it would be valuable to investigate whether similar effects occur in birds by testing the impact of NO inhibition on spatial task performance.

In addition to its role in learning, NO has been linked to aging-related cognitive decline (Domek-Łopacińska & Strosznajder, 2010; Law et al., 2001; McCann, 1997). In mammals, a reduction in NO levels has been associated with impaired synaptic plasticity and decreased cognitive function (Domek-Łopacińska & Strosznajder, 2010). Rodent studies have shown that aged rats exhibit a reduced number of NADPH-d-positive neurons in the cerebral cortex and striatum, as well as reduced NOS activity in the cerebellum. These age-related neurochemical changes were associated with impairments in radial arm maze performance, suggesting a decline in NO-mediated mechanisms of spatial learning (Yamada et al., 1996). Although little is known about NO’s role in aging birds, evidence from zebra finches reveals that NO expression decreases in song nuclei with age, affecting song plasticity (Wallhäusser-Franke et al., 1995). Given pigeons’ long lifespan and sustained cognitive abilities, investigating NO’s potential contribution to age-related memory decline could provide insights into its broader neuromodulatory functions.

### Evolutionary and Comparative Perspectives

The widespread presence of NADPH-d across avian brain regions indicates that NO plays a general modulatory role in sensory and cognitive processing, with species-specific variations likely reflecting ecological adaptations. Studies in quails, chickens, and pigeons have demonstrated a broadly conserved NO distribution across forebrain, midbrain, and hindbrain structures (Atoji et al., 2001; Brüning, 1993; Panzica et al., 1994). However, distinct ecological and behavioral specializations may influence NO expression in key regions. For example, species-specific differences have been observed, such as stronger NO expression in the pigeon hippocampus and olfactory bulb compared to non-navigating species (Brüning, 1993; Panzica et al., 1994). This suggests a potential link between NO signaling and spatial cognition, where pigeons, known for their exceptional navigational abilities (Gagliardo et al., 1999; Rook et al., 2023), might rely on NO-mediated plasticity to process and adapt spatial information. Supporting this notion, rodent studies have shown that NO regulates hippocampal plasticity and spatial memory, with NO inhibition leading to impairments in learning tasks, such as the Morris water maze (Harooni et al., 2009; Majlessi et al., 2008). Similarly, food-storing birds exhibit elevated NO expression in the hippocampus, likely supporting the greater spatial memory demands required for efficient retrieval of cached food (Krebs et al., 1996).

Conversely, in species that rely heavily on vocal communication, such as songbirds and parrots, NO expression is particularly elevated in auditory-processing regions (Brauth et al., 2005; Cozzi et al., 1997; Wallhäusser-Franke et al., 1995). In vocal learners, NO has been detected in song-control nuclei, including HVC, RA, and Area X, with species-specific variations suggesting adaptations to distinct vocal learning strategies (Brauth et al., 2005; Cozzi et al., 1997). Among these species, budgerigars exhibit a higher density of NADPH-d-positive neurons in the anterior forebrain, particularly in the nucleus basalis (Bas), a region associated with vocal learning and auditory processing (Cozzi et al., 1997). In contrast, zebra finches show an age-dependent decline in NADPH-d expression within song nuclei, potentially linked to differences between lifelong learners (e.g., budgerigars) and species with restricted vocal learning periods (e.g., zebra finches) (Brauth et al., 2005; Wallhäusser-Franke et al., 1995). These findings support the idea that NO expression in song-related circuits may be shaped by species-specific vocal learning demands and plasticity mechanisms.

In pigeons, NO activity was detected in the arcopallium and entopallium, brain regions associated with motor control and visual processing (Atoji et al., 2001). However, NO expression in these areas appeared weaker than previously reported by Atoji et al. (2001), raising questions about potential methodological differences or individual variability. While the functional significance of this variation remains uncertain, comparative analyses suggest that NO expression patterns across species may be closely linked to distinct sensory, motor, and cognitive specializations.

Future research should explore whether NO-mediated memory flexibility is shaped by ecological pressures, particularly in species with higher cognitive plasticity. If NO plays a key role in adaptive forgetting, comparative studies across avian taxa could reveal whether this mechanism is more pronounced in species requiring high behavioral flexibility. Further experimental work involving NO manipulation during extinction learning paradigms may provide deeper insights into how NO contributes to memory regulation and adaptive decision-making across vertebrates.

## Conclusion

This study provides anatomical evidence that NO contributes to dopaminergic modulation in the pigeon NCL, potentially supporting mechanisms that balance memory retention and forgetting—mirroring findings in Drosophila. Its presence in the hippocampus and links to aging further underscore its broader relevance to avian cognition. Behavioral studies are now needed to test whether NO influences memory persistence and cognitive flexibility in birds. By establishing this neuroanatomical framework, our findings advance the understanding of NO as a conserved neuromodulatory factor in memory processing across species.

## Acknowledgement

Supported by the Deutsche Forschungsgemeinschaft (DFG), Project-ID 316803389 – SFB 1280, and SFB1372 Project-ID 395940726.

## Author contributions

AS: conceptualization, data curation, data analysis, writing (original draft), visualization; MZ: data curation, visualization; KH: data curation; data analysis, OG: conceptualization, writing (editing), funding acquisition, resources; NR: conceptualization, data curation, data analysis, visualization, writing (editing), supervision.

## Data availability

Data will be made available upon request from the corresponding author

